# Nucleomorph phylogenomics suggests a deep and ancient origin of cryptophyte plastids within Rhodophyta

**DOI:** 10.1101/2024.03.10.584144

**Authors:** Lukas V. F. Novak, Sergio A. Muñoz-Gómez, Fabian van Beveren, Maria Ciobanu, Laura Eme, Purificación López-García, David Moreira

**Affiliations:** Unite d’Ecologie Systematique et Evolution, CNRS, AgroParisTech, Universite Paris-Saclay, Gif-sur-Yvette, France; Univ Brest, CNRS, Ifremer, EMR 6002 BIOMEX, IRP 1211 MicrobSea, Unite Biologie et Ecologie des Ecosystemes marins Profonds BEEP, F-29280 Plouzane, France; Department of Biological Sciences, Purdue University, IN, USA; Laboratory of Microbiology, Wageningen University & Research, Netherlands

**Author notes:** (LVFN), (DM).

**Keywords:** evolution, symbiosis, algae, protists, plastids, nucleomorphs, Rhodophyta, Cryptophyta

## Abstract

The evolutionary origin of red algae-derived complex plastids remains obscure. Cryptophyta, one of four eukaryotic lineages harboring these plastids, still contain nucleomorphs, which are highly reduced remnants of red algal nuclei. The genes present on nucleomorph genomes can be used for phylogenomic reconstruction in order to unravel the evolutionary origin of all red complex plastids and provide data independent from previously analyzed plastid-encoded proteins. Here, we leverage these genes in a first attempt at pinpointing the position of cryptophyte nucleomorphs within a comprehensive diversity of Rhodophyta, including new sequence representatives from seven deep-branching red algae. Our analysis, supported by a series of rigorous topology tests, places cryptophyte nucleomorphs as sister to the extremophilic, freshwater subphylum Cyanidiophytina. This conflicts with previously published analyses based on plastidial genes that placed red complex plastids closer to the mesophilic Rhodophytina. Regardless of exact placement, our results reject a nucleomorph origin within any known subgroup of Rhodophyta, instead suggesting an ancient origin of complex red plastids among the deepest branches of the red algal tree of life.

## Introduction

Complex plastids are typically photosynthetic organelles found in multiple unrelated groups of eukaryotes, derived from a symbiosis with another photosynthetic eukaryote, as opposed to primary plastids derived directly from a symbiosis with a cyanobacterium (Irisarri *et al*., 2022; Füssy & Oborník, 2024; Novák Vanclová & Dorrell, 2024). The origin of complex plastids of red algal (Rhodophyta) ancestry found in photosynthetic members of the Cryptista, Alveolata, Stramenopiles, and Haptista (“CASH”) remains one of the great unresolved questions of eukaryotic evolution. Two competing hypotheses have tried to explain the presence of these organelles of apparently shared origin in these four distantly related lineages of hosts (Yoon *et al*., 2002; Burki *et al*., 2020). On the one hand, the chromalveolate hypothesis (Cavalier-Smith, 1999) posits a single origin of these plastids from a single secondary endosymbiosis event in a common ancestor of CASH, and its subsequent vertical inheritance accompanied by multiple independent plastid losses in non-photosynthetic CASH species. On the other hand, the serial endosymbiosis hypothesis (Sanchez-Puerta & Delwiche, 2008) postulates that secondary endosymbiosis occurred more recently in a single CASH lineage, after the four lineages had already diverged, and spread to the rest horizontally *via* tertiary, or even higher-degree, endosymbioses. This serial endosymbiosis hypothesis has been proposed in multiple versions positing different numbers and order of symbiotic events (Sanchez-Puerta & Delwiche, 2008; Petersen *et al*., 2014; Stiller *et al*., 2014; Bodył, 2018).

Regardless of which particular scenario is true, the identity of the red algal donor of complex plastids also remains uncertain. Phylogenomic analyses of plastid-encoded genes (Yoon *et al*., 2002; Iida *et al*., 2007; Janouškovec *et al*., 2010; Stiller *et al*., 2014; Ševčíková *et al*., 2015; Kim *et al*., 2017) have consistently placed the CASH complex plastids in the vicinity of the Rhodophytina, the mesophilic branch of Rhodophyta, as opposed to the ancestrally extremophilic, freshwater Cyanidiophytina (Yoon *et al*., 2017). However, our insufficient understanding of phylogenetic relationships among red algal groups, as well as poor taxon sampling of rhodophyte genomes, have for a long time limited the usefulness of such studies in precisely pinpointing the donor taxon. Recent large-scale phylogenomic analyses of plastid- and mitochondrion-encoded genes (Muñoz-Gómez *et al*., 2017; van Beveren *et al*., 2022) have greatly advanced our understanding of the internal phylogeny of the phylum Rhodophyta. These analyses support the basic division between Cyanidiophytina (class Cyanidiophyceae) and Rhodophytina, and split the latter into two monophyletic subphyla: Proteorhodophytina (classes Compsopogonophyceae, Porphyridiophyceae, Rhodellophyceae, and Stylonematophyceae), comprising red algae with single-celled or simple multicellular body plans, and Eurhodophytina (classes Bangiophyceae and Florideophyceae), with complex multicellularity.

To attempt to phylogenetically place the CASH plastids within the tree of Rhodophyta, we used an alternative source of genetic data – the nucleomorph genomes of Cryptophyta (Cryptista). Nucleomorphs (NMs) are highly reduced nuclei of archaeplastid algae that reside in the periplastidial space of certain complex plastids. There are at least three independent examples of NMs derived from green algae (Ludwig & Gibbs, 1989; Sarai *et al*., 2020), but only one derived from red algae (Ludwig & Gibbs, 1987; Douglas *et al*., 2001). Cryptophyte NMs are relatively well studied; phylogenetic analyses of their rRNAs have suggested a rhodophyte origin (Douglas *et al*., 1991; Maier *et al*., 1991), and they have even been included in explorations of rhodophyte genomic diversity (Wong *et al*., 2023). However, to our knowledge, there has not been any published phylogenomic analysis attempting to locate the position of cryptophyte NMs within Rhodophyta, with a single exception where NMs were not the focus and their placement was not discussed (Strassert *et al*., 2021).

Cryptophyte NMs contain several hundred open reading frames (ORFs) that can be divided into three categories: conserved ORFs with homologues outside cryptophyte NMs, nucleomorph-specific ORFs or “nORFs” with homologues only in other cryptophyte NMs, and “ORFans” without recognizable sequence similarity to any other known sequence (Moore *et al*., 2012). Only conserved ORFs are therefore useful for phylogenomic analyses aimed at placing NMs in a broader eukaryotic context. With this goal in mind, we have improved the taxon sampling of red algae by sequencing seven novel nuclear genomes and performed a series of phylogenomic investigations into the position of cryptophyte NMs among Rhodophyta using a large dataset of 180 conserved ORFs. Our analyses reject the monophyly of cryptophyte NMs with any single class of mesophilic rhodophytes and support their position as sister to Cyanidiophytina.

## Materials and Methods

To improve the taxon sampling of nucleus-encoded protein sequences of Rhodophyta available for phylogenomic analyses, we sequenced DNA of seven red algae: *Galdieria* sp. ACUF 613 (Cyanidiophyceae), *Madagascaria erythrocladioides* (Compsopogonophyceae), *Erythrolobus coxiae*, *Porphyridium aerugineum*, *Timspurckia oligopyrenoides* (Porphyridiophyceae), *Dixoniella grisea*, and *Rhodella violacea* (Rhodellophyceae). For culturing, DNA extraction, and sequencing methods see (van Beveren *et al*., 2022). Raw genomic reads for *D. grisea*, *E. coxiae*, *Galdieria* sp. ACUF 613, *M. erythrocladioides*, *P. aerugineum*, and *T. oligopyrenoides* were obtained by Illumina HiSeq 2500 and NovaSeq 6000. Raw genomic reads for *R. violacea* were obtained by Illumina HiSeq 2500 and NovaSeq 6000 plus Oxford Nanopore MinION Mk 1B. The reads were assembled using SPAdes assembler, version 3.14.1 (Prjibelski *et al*., 2020). Nuclear contigs were separated from organellar and prokaryotic contamination-derived sequences using Whokaryote, version 1.1.2 (Pronk & Medema, 2022). Raw reads were mapped to resulting contigs using the BWA-MEM (Li, 2013) algorithm (github.com/lh3/bwa), and BLASTn (Altschul *et al*., 1990) was used to search the contigs against the NCBI nt database, release 240. Results of the read mapping and BLASTn search were visualized in BlobTools2 (Challis *et al*., 2020) and used for additional manual decontamination, resulting in preliminary genomic assemblies (not shown).

Previously published transcriptomic reads for *M. erythrocladioides* (SRX554333), *T. oligopyrenoides* (SRX554192), and *R. violacea* (ERX2100208) were downloaded from the NCBI SRA database (Sayers *et al*., 2021) and mapped to their respective genomic assemblies using the BWA-MEM algorithm and visualized in the Integrative Genomics Viewer, version 2.9.4 (Robinson *et al*., 2011). Sets of >200 protein-coding gene models were manually predicted for each of the three assemblies with mapped transcripts and used as training sets for the online AUGUSTUS gene predictor (Hoff & Stanke, 2013). The resulting species parameters, as well as previously available *Galdieria sulphuraria* (Cyanidiophyceae) species parameters, were used in parallel for AUGUSTUS automatic gene prediction based on each of the seven genomic assemblies. The completeness of the individual sets of AUGUSTUS-predicted genes was evaluated using BUSCO, version 5.4.3, with the eukaryota_odb10 dataset (Manni *et al*., 2021) and the one with the highest estimated completeness was selected for each of the seven assemblies. These preliminary predicted proteomes were searched for protein sequences of interest for use in further analyses.

In order to maximize the independence of our phylogenomic matrix from plastid data, we started with a seed dataset of 228 NM-encoded proteins (markers) from the non-photosynthetic cryptophyte *Cryptomonas paramecium*, which were annotated as not being associated with the plastid or photosynthesis (Tanifuji *et al*., 2010). The 228 *C. paramecium* NM-encoded protein sequences from the “Conserved ORFs” dataset (Tanifuji *et al*., 2010) were searched against a custom protein database consisting of the seven novel rhodophyte preliminary predicted proteomes plus three cryptophyte NM predicted proteomes downloaded from the NCBI GenBank database, release 245, and multiple predicted proteomes representing a broad diversity of eukaryotes downloaded from the EukProt database, version 2 (Richter *et al*., 2022). Two best (lowest E-value) BLASTp hits were recovered per proteome for each query sequence. The sequences were aligned using MAFFT, version 7.455 (Katoh & Standley, 2013) and resulting alignments were trimmed using BMGE, version 1.12 (Criscuolo & Gribaldo, 2010) and then used to reconstruct initial single-gene trees using FastTree, version 2.1.8 with the JTT+CAT model (Price *et al*., 2010). The trees (supplementary file S1) were manually inspected and used to: 1) select 180 proteins that produced trees where all the NM sequences branched together to the exclusion of all other sequences (no clear alternative patterns were observed in the trees of the discarded proteins), 2) select 54 proteomes most relevant to the studied question, i.e. four from cryptophyte NMs and 50 from four lineages of Archaeplastida (Rhodophyta, *Rhodelphis*, Glaucophyta, and Viridiplantae), and 3) when paralogs were present, select a single paralog per tree – determined by a shorter branch. This selection resulted in an initial phylogenomic dataset of 180 markers in 54 species. Details about the selected species are shown in supplementary file S2.

The selected protein sequences were gathered in full length into 180 single-gene multi-FASTA files. PREQUAL, version 1.01 (Whelan *et al*., 2018) was used to identify and mask positions with non-homologous characters and the masked sequences were aligned using MAFFT, version 7.455 (Katoh & Standley, 2013). The alignments were then trimmed using DIVVIER (-partial -mincol 28), version 1.01 (Ali *et al*., 2019). The resulting alignments (supplementary file S3) were used to reconstruct single-marker phylogenetic trees as well as to assemble a concatenated supermatrix using phyutility, version 2.2 (Smith & Dunn, 2008). Details about the matrix are shown in supplementary file S4.

The supermatrix (supplementary file S5) was analyzed with IQ-TREE, version 2.2.0 (Minh *et al*., 2020) using the substitution model LG+C60+G+F to account for across-site compositional heterogeneity (Si Quang *et al*., 2008) and both ultrafast bootstrap (-bb 1000) and nonparametric bootstrap (-bo 100) for the maximum likelihood (ML) phylogenetic analysis, and with PhyloBayes-MPI, version 1.8c (Lartillot *et al*., 2013) with the CAT-GTR substitution model (Lartillot & Philippe, 2004) for the Bayesian inference phylogenetic analysis (figure 1). The PhyloBayes-MPI analysis with two chains was run for more than 50,000 generations. The chains did not converge (maxdiff=1, meandiff=0.0190534, burn-in: 1000, sub-sampling every 10 trees), as is often the case with large phylogenomic datasets. Trace files were inspected with Tracer (Rambaut *et al*., 2018). Trees were visualized and edited using TreeViewer, version 2.1.0 (Bianchini & Sánchez-Baracaldo, 2024).

**Figure 1.**
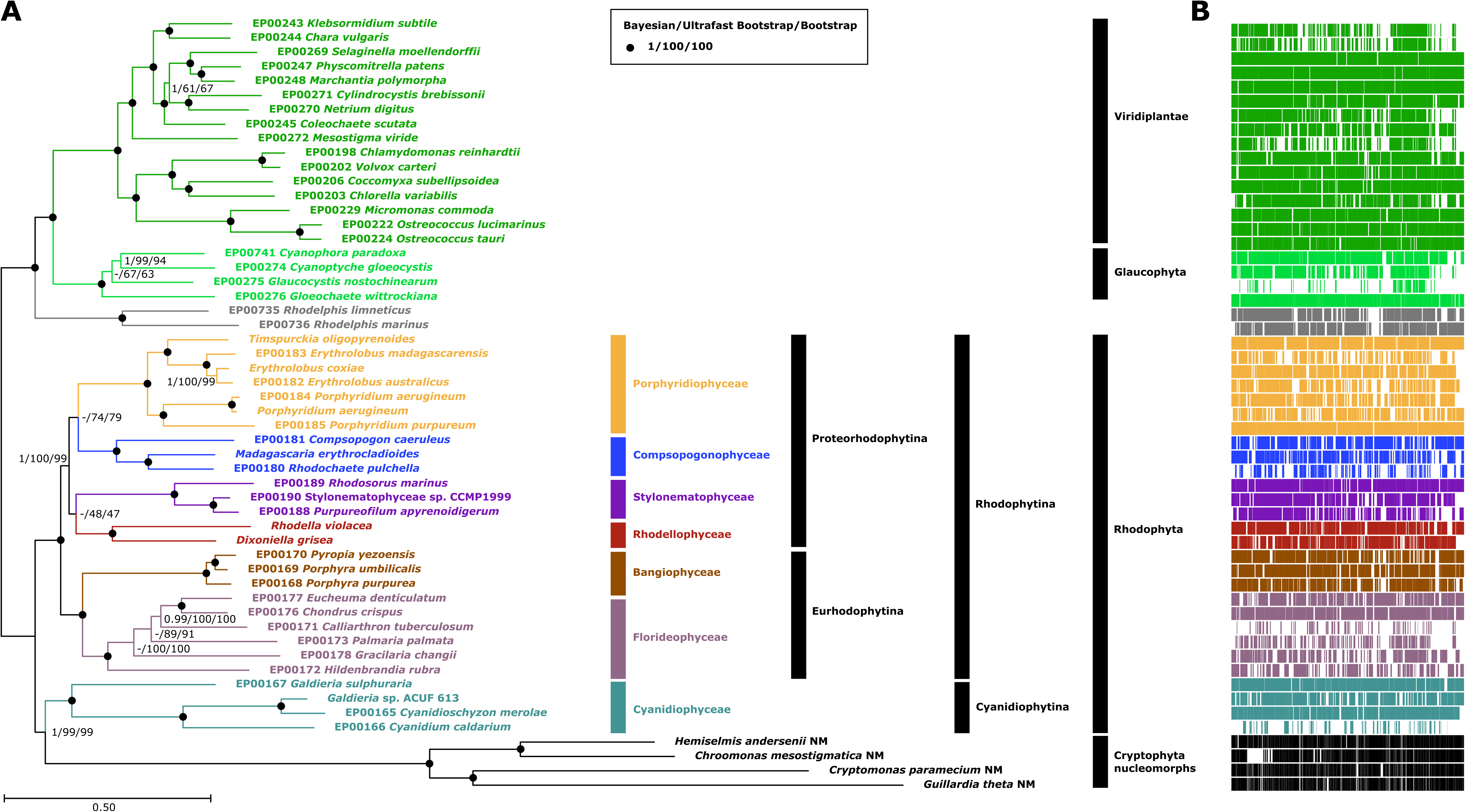
Phylogenetic tree placing cryptophyte NMs among Archaeplastida. (**A**) Maximum likelihood (LG+C60+G+F) phylogenetic tree constructed from a concatenation of 180 nucleus/nucleomorph-encoded proteins (51,610 positions, 54 taxa). Branch support values are derived from Bayesian posterior probability (CAT-GTR) and maximum likelihood ultrafast and nonparametric bootstrap. (**B**) Completeness of the individual datasets shown as a schematic representation of the complete supermatrix alignment (gaps in white). The conspicuous block of missing data in *C. mesostigmatica* is related to the previously reported lineage-specific loss of proteasome-related genes in this genome (Moore *et al*., 2012).

Single-marker trees (supplementary file S6) were reconstructed using IQ-TREE (-bb 1000 -m LG+C60+G+F) version 2.2.0. PhyKIT, version 1.11.3 (Steenwyk *et al*., 2021) was used to extract branch lengths in the 180 single-marker trees and this information guided further tests (see below) of topology robustness by sequential removal of markers or individual sequences.

In test 1, all markers were sorted by the length of the root branch of cryptophyte NMs (first common ancestor to last common ancestor of NMs) from longest to shortest to generate additional supermatrices, by removing, in a stepwise fashion, 10% of the markers. In test 2, all markers were sorted by the average compound length of cryptophyte NM branches (first common ancestor of NMs to tips) from longest to shortest to generate 8 supermatrices, by removing, in a stepwise fashion, 10% of the markers. In test 3, only individual sequences were removed, not entire markers. All NM sequences were sorted by their compound branch length (first common ancestor of NMs to tips) from longest to shortest to generate 8 supermatrices, by removing, in a stepwise fashion, 10% of the NM sequences. In this test, 14 sequences (from NMs, Rhodophyta, and *Rhodelphis*), identified in the IQ-TREE-generated single-marker trees as disrupting the NM monophyly, were removed together with the first 10% longest-branch sequences. Details about the removed markers and sequences and their sorting in tests 1–3 are shown in supplementary file S7.

In test 4, IQ-TREE, version 2.2.0 was used to calculate site-specific evolutionary rates (--rate-mlrate -m LG+C60+G+F) for each position in the original supermatrix (supplementary file S8) and SiteStripper, version 1.02 (github.com/hverbruggen/SiteStripper) was used to generate 8 supermatrices, each by removing, in a stepwise fashion, 10% of the positions with the highest evolutionary rates.

In test 5, amino acid compositional bias of the species in the supermatrix was determined according to the χ2 metric as described before (Viklund *et al*., 2012; Martijn *et al*., 2018; Muñoz-Gómez *et al*., 2019). Occurrences of all amino acids in all positions in the matrix were counted and the positions were sorted according to the ratio F+S+I+N+K / A+P+V+D+M in NMs divided by the ratio F+S+I+N+K / A+P+V+D+M in all other species. In order to avoid dividing by zero, an arbitrary value of 0.01 was added to all the amino acid occurrence values before the calculation. Details about this calculation and sorting are given in supplementary file S9. A custom Python script was used to generate 8 supermatrices, each with 10% of the remaining positions removed starting with those with NMs most enriched in F+S+I+N+K and other species most depleted in A+P+V+D+M.

The modified supermatrices generated in tests 1–5 were used to reconstruct ML trees (IQ-TREE, -bb 1000 -m LG+C60+G+F) and the ultrafast bootstrap values supporting the original or alternative positions of NMs within the resulting topologies were plotted in figure 2.

**Figure 2.**
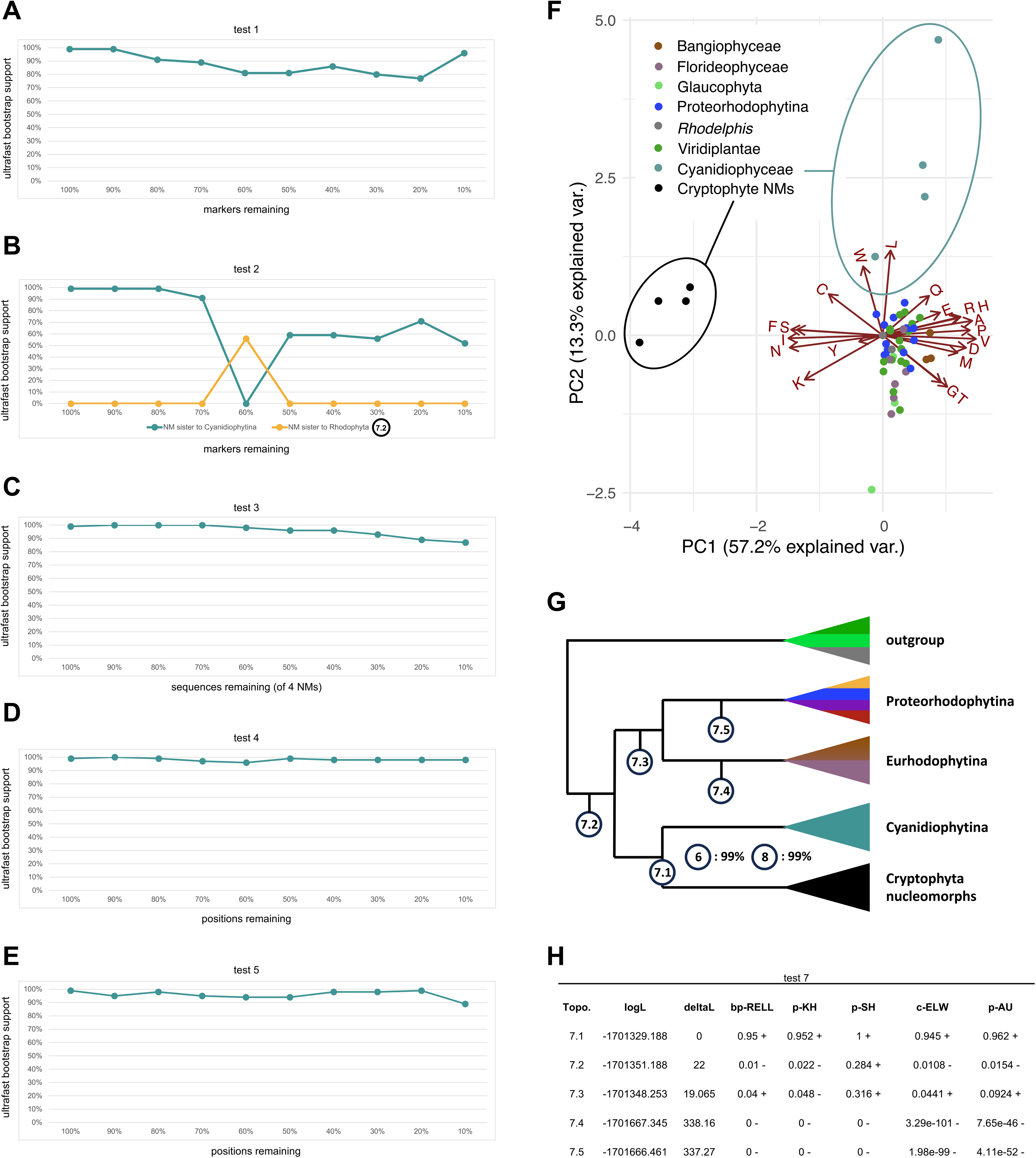
Results of the topology robustness tests 1 to 8. (**A–E**) Changes in ultrafast bootstrap support for the sister relationship between cryptophyte NMs and Cyanidiophytina or alternative topologies as calculated based on reduced supermatrices in tests 1–5. (**F**) Principal component analysis of the taxa used in the phylogenetic analyses based on the amino acid composition of their sequences. (**G**) Simplified schematic representation of the alternative placements of cryptophyte NMs used in test 7 and ultrafast bootstrap support for the sister relationship between cryptophyte NMs and Cyanidiophytina as calculated based on modified supermatrices in tests 6 and 8. (**H**) Results of approximately unbiased topology tests (test 7).

To further evaluate the effects of compositional heterogeneity on phylogenetic inference, in test 6, we recoded the supermatrix into a four-character state amino acid alphabet that minimizes the compositional heterogeneity present in the data. The program minmax-chisq, which implements the methods of Susko and Roger (Susko & Roger, 2007), was used to find the best recoding scheme for our supermatrix. The best recoding scheme (S4) was found to be NQFPV, RDCLKMST, AEGHIY, and W. Phylogenetic analyses on the recoded supermatrix were performed with IQ-TREE, version 2.2.2.7 and the across-site compositionally heterogeneous GTR+C60(S4)+F+R6 model. C60(S4) is an adaptation of the C60 mixture model to four-character states. It is obtained by adding the frequencies of the amino acids that belong to each bin in the dataset-specific four-character state scheme S4.

In test 7, we calculated ML trees (IQ-TREE, -bb 1000 -m LG+C60+G+F) with alternative topologies by constraining possible relationships under following scenarios: 1) unconstrained original tree, i.e. NMs+Cyanidiophytina, 2) NMs outside Rhodophyta, 2) NMs+Rhodophytina, 4) NMs+Eurhodophytina, and 5) NMs+Proteorhodophytina. AU-test (Shimodaira, 2002) was used as implemented in IQ-TREE (-m LG+C60+G+F -n 0 -zb 1000 - au) to test the alternative hypotheses.

To evaluate the effect of functional divergence in the protein sequences, in test 8 we used the software FunDi, version 1.1 (Gaston *et al*., 2011) to estimate functionally divergent sites in the branch that separates nucleomorphs from all other nuclear genomes. About 7% of all amino acid sites were estimated to be functionally divergent with a posterior probability of > 0.5. These sites were subsequently removed using a custom Perl script in order to reduce potential artefacts from model misspecification, since current phylogenetic models do not properly capture functional divergence in proteins (Inagaki *et al*., 2003; Muñoz-Gómez *et al*., 2022). The resulting supermatrix consisted of 47,951 amino acids. Phylogenetic analyses of this supermatrix without functionally divergent sites were performed with IQ-TREE version 2.2.2.7 and the across-site compositionally heterogeneous LG+C60+G4+F mixture model.

In order to test the influence of taxon sampling on the resulting topologies, as well as to incorporate new and relevant genomic assemblies recently published, we performed tests 9 and 10 with varying number of included taxa. To construct datasets for these tests, we searched the newly added assemblies using HMMER, version 3.4 (Eddy, 2011) with HMM profiles constructed from the original 180 protein alignments. The best scoring results of the HMMER searches were included in a set of 180 datasets used to reconstruct initial IQ-TREE (LG+C60+G+F, -bb 1000) phylogenetic trees (supplementary file S11) and to remove sequences forming exceptionally long branches in a process mirroring the creation of the initial phylogenomic dataset.

In test 9 we included five new cryptophyte NM genomic assemblies (Kim *et al*., 2022; George *et al*., 2023) and repeated the initial ML phylogenetic tree reconstruction in order to determine the relative branch lengths of all nine NMs (first common ancestor of NMs to tips). Then we constructed eight additional supermatrices by removing, in a stepwise fashion, individual NM datasets from the analyses in order of decreasing branch length. This was intended to evaluate the influence of the overall NM branch length on the topology.

In test 10 we evaluated the influence of the outgroup composition by including 68 additional taxa from the EukProt database (Richter *et al*., 2022) covering a broad diversity of eukaryotes beyond the Archaeplastida and repeating the initial phylogenetic analysis to sort all the outgroup taxa by their relative branch length (first common ancestor of Rhodophyta+NMs to tips). Eight additional supermatrices were generated by removing, in a stepwise fashion, 10% of the outgroup taxa with the longest branches.

All alternative trees used in the tests can be found in supplementary file S10. All bioinformatic methods are summarized in supplementary file S12.

## Results

We sequenced the genomes of seven red algae, which, together with previously published genome- and transcriptome-derived predicted proteomes available in the EukProt database (Richter *et al*., 2022), cover all seven classes of Rhodophyta and, for the first time, allow analyzing the cryptophyte NMs phylogenetic position in relation to the full known diversity of red algae. Our final phylogenomic sequence supermatrix contained 51,610 amino acid positions belonging to 180 conserved protein markers (see annotations in supplementary file S4); it included 54 species covering Viridiplantae, Glaucophyta, *Rhodelphis*, all seven classes of Rhodophyta (including three of the four known orders of Cyanidiophyceae (Park *et al*., 2023)), and NMs of four distantly related cryptophyte species: *C. paramecium*, *Guillardia theta*, *Chroomonas mesostigmatica*, and *Hemiselmis andersenii* (Hoef-Emden *et al*., 2002; Kim & Archibald, 2013; Greenwold *et al*., 2023).

We carried out phylogenomic analyses of the sequence matrix by maximum likelihood (ML) with the LG+C60+G+F mixture model and by Bayesian inference with the CAT-GTR model. The resulting trees were outgroup-rooted on the non-rhodophyte Archaeplastida. Both analyses (figure 1) recovered the expected topology of Archaeplastida and Rhodophyta, including the monophyly of the only recently recognized Proteorhodophytina (Muñoz-Gómez *et al*., 2017), with full support (1/100/100: Bayesian posterior probability/ultrafast bootstrap/nonparametric bootstrap). Surprisingly, the analyses recovered the cryptophyte NMs as sister to Cyanidiophytina with strong support (1/99/99). This result is conflicting with all previously published phylogenomic analyses of CASH plastids (including Cryptophyta) based on plastid-encoded genes, which unanimously placed CASH as sister to the other major branch of red algae, the Rhodophytina.

We tested the robustness of the recovered topology in a series of analyses (tests 1–8, see Materials and Methods) designed to uncover potential influence of long-branch attraction or sequence compositional bias on the ML results (figure 2). If the recovered topology was a result of these phylogenetic artefacts, the progressive removal of divergent data (individual positions or marker genes) from the alignment would likely result in a clear trend of lowering statistical support and possibly towards an alternative topology.

Among tests 1–5, where we inferred ML trees from supermatrices generated by gradually removing divergent data in increments of 10% (figure 2A–E), only in test 2 (sequential removal of markers by the average compound length of cryptophyte NM branches), we observed a substantial drop of the support for the monophyly of cryptophyte NMs and Cyanidiophytina. An alternative topology (NMs sister to all Rhodophyta) rose over 50% support in a single tree (with 60% of markers remaining). None of the other tests showed this alternative topology and the support for NMs sister to Cyanidiophytina always remained above 90%.

The ML tree from test 6, where we recoded the amino acid supermatrix into a four-character state alphabet, presented the same topology as the original one with 99% support for NMs as sister group to Cyanidiophytina (figure 2G). The AU test (test 7, figure 2H) rejected all topologies except the original ML tree (figure 2G: topology 7.1). The topology consistent with the previous plastid-encoded proteins analyses, i.e., NMs sister to Rhodophytina, could not be rejected, although the p-value (0.0924) was on the cusp of the cut-off for significance (figure 2G: topology 7.3). Notably, the topology briefly supported in test 2 (figure 2G: topology 7.2) was rejected by the AU test (p-value=0.0154). The ML tree reconstructed based on a modified supermatrix with 7% of the most functionally divergent sites removed (test 8) again recovered the original topology with 99% support for the monophyly of NMs and Cyanidiophytina (figure 2G).

After inclusion of five new NM assemblies from the genus *Cryptomonas* (Kim *et al*., 2022; George *et al*., 2023), the ultrafast bootstrap support for cryptophyte NMs being sister to Cyanidiophytina dropped only slightly to 94% (figure 3A) and remained above 90% throughout test 9, where we sequentially removed NMs from the longest–branching one to the shortest-branching one. Only when two NMs remained in the dataset, did the support drop to 78%, and with only a single NM present, an alternative topology (NMs sister to Rhodophytina; 7.3 in figure 2G) was slightly favored with an ultrafast bootstrap value of 52% (figure 3C).

**Figure 3.**
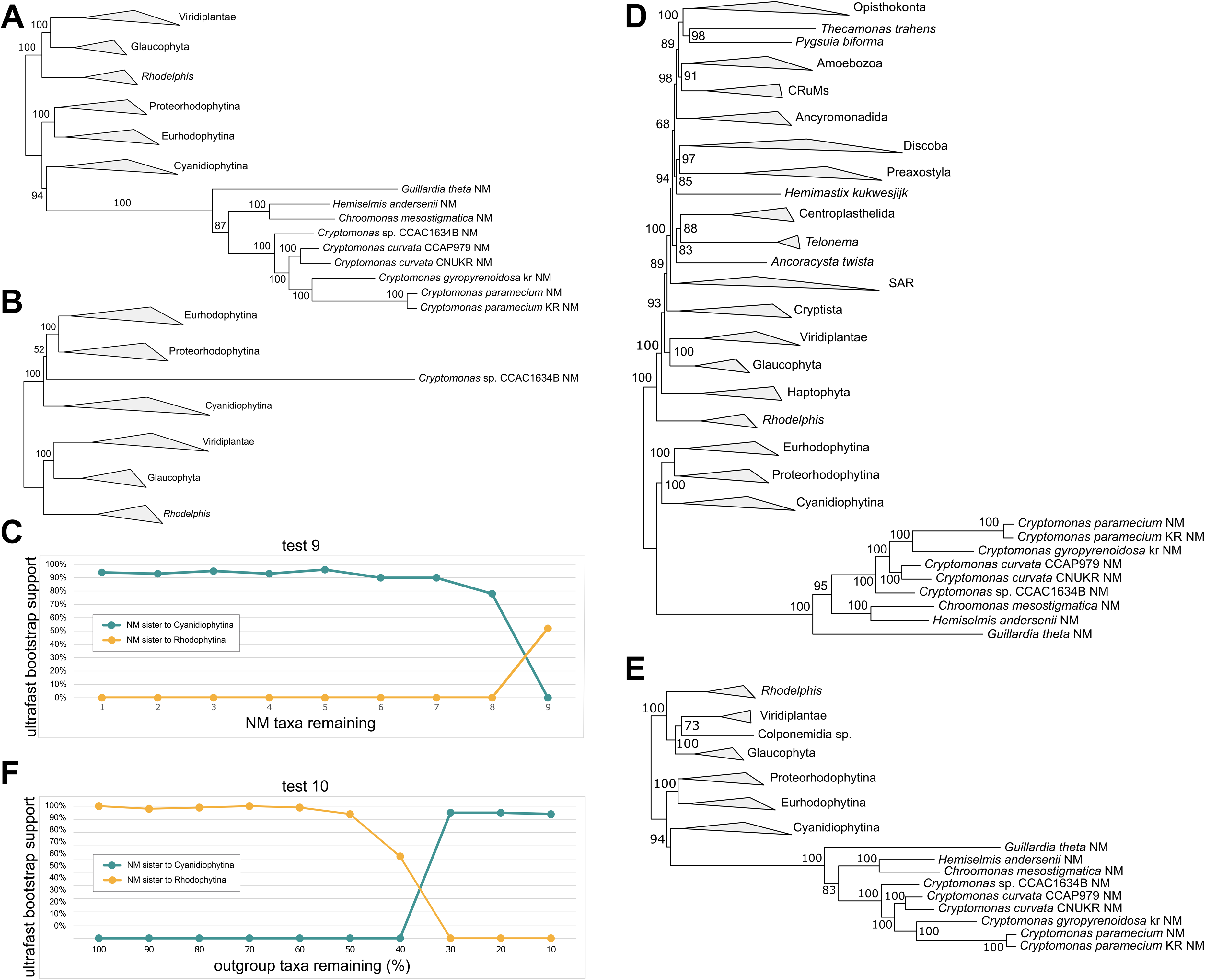
Results of the topology robustness tests 9 and 10 after inclusion of additional taxa. (**A**) Maximum likelihood (LG+C60+G+F, 1000x ultrafast bootstrap) tree including all 9 available NM genomic datasets (maximum number of NMs in test 9). (**B**) Maximum likelihood (LG+C60+G+F, 1000x ultrafast bootstrap) tree including only the single NM genomic dataset with the shortest branch (minimum number of NMs in test 9). (**C**) Changes in ultrafast bootstrap support for the sister relationship between cryptophyte NMs and Cyanidiophytina or alternative topologies as calculated based on the progressive removal of cryptophyte NMs (test 9). (**D**) Maximum likelihood (LG+C60+G+F, 1000x ultrafast bootstrap) tree including all 9 available NM genomic datasets and 68 additional outgroup taxa (maximum number of taxa in test 10). (**E**) Maximum likelihood (LG+C60+G+F, 1000x ultrafast bootstrap) tree including all 9 available NM genomic datasets and only 10% of the outgroup taxa with shortest branches (minimum number of taxa in test 10). (**F**) Changes in ultrafast bootstrap support for the sister relationship between cryptophyte NMs and Cyanidiophytina or alternative topologies as calculated based on increasingly reduced numbers of taxa (test 10).

After inclusion of 68 additional outgroup taxa from a broad diversity of eukaryotes, the cryptophyte NMs branched sister to all Rhodophyta (7.2 in figure 2G) and the tree was not fully consistent with the expected eukaryotic phylogeny (e.g. Haptophyta branched within Archaeplastida). When 50% of the outgroup taxa with longest branches were removed in test 10, the support for the alternative topology started to diminish and at 30%, the originally recovered topology of NMs sister to Cyanidiophytina received support of 95% and remained over 90% even at 20% and 10% of remaining outgroup taxa (figure 3E, 3F).

## Discussion

Our analyses show that the hypothesis of cryptophyte NMs branching sister to Cyanidiophytina is the best supported one by the phylogenetic analyses of conserved genes still preserved in the cryptophyte nucleomorph genomes. There is no strong indication that this result was influenced by long-branch attraction or sequence compositional bias. In fact, reanalyzing the supermatrix with fewer divergent data did not show any trend towards alternative topologies. Interestingly, this position was also recovered in a previous pan-eukaryotic phylogenomic analysis which included NM data (Strassert *et al*., 2021). The alternative position of NMs as sister to Rhodophytina is not strongly rejected by the NM data and, given its recovery in previous plastid genome phylogenetic analyses (Iida *et al*., 2007; Janouškovec *et al*., 2010; Ševčíková *et al*., 2015; Kim *et al*., 2017), we cannot at this stage favor one over the other.

It is difficult to imagine a mechanism by which a red algal plastid and a red algal nucleus of different origins would end up together as components of the current complex plastid of cryptophytes. Therefore, we should assume that they indeed derive from the same red algal lineage and their different phylogenetic signals are not reflecting different evolutionary histories. The discrepancy between the two topologies, 1) NMs sister to Rhodophytina, well supported by plastidial data and 2) NMs sister to Cyanidiophytina, well supported by NM data, can be explained either by an undiscovered phylogenetic artefact, or by an unknown gene transfer event that resulted in different phylogenetic signal in the two sources of genomic data.

The plastidial and NM genomes presumably undergo different evolutionary pressures, and it is conceivable that this resulted in a so far undiscovered systematic compositional bias in one or both sets of genomes, leading to different phylogenetic signals. Alternatively, either the plastidial or NM genomes might be enriched in genes of a different origin as a result of ancient horizontal gene transfer (HGT). For example, if the entire complex plastid was derived from a Rhodophytina-related ancestor, as suggested by the plastidial data, the Cyanidiophytina-like phylogenetic signal could be a result of predation on Cyanidiophytina prey by the cryptophyte host and frequent HGT replacement of genes in the NM genome.

An opposite scenario could be also conceived, in which plastidial genes were replaced by HGT from Rhodophytina prey. This would be somewhat analogous to the “red carpet” hypothesis explaining the presence of rhodophyte proteins in proteomes of species with secondary plastids of green origin (Ponce-Toledo *et al*., 2019). A period of retention of two distinct endosymbionts, as exemplified by certain dinotoms (Hehenberger *et al*., 2014), or an even more complicated endosymbiotic architectures, like the karyoklepty described in ciliates (Johnson *et al*., 2007), could also have happened in an ancestor of cryptophytes. Such interactions could have resulted in HGT into one of the symbiotic genomes. Such scenarios are highly speculative and very difficult to test, given the close relationship between the two alternative source lineages and the challenges of precisely reconstructing single-gene phylogenetic trees.

Most importantly, our analyses neither show a close relationship between cryptophyte NMs and any of the six classes of Rhodophytina, nor their origin from within the known extant diversity of Cyanidiophytina. On the contrary, they support a deep and ancient origin of the cryptophyte NMs and, therefore, presumably also of all CASH plastids, within red algae. They likely emerged as sister to either Cyanidiophytina or Rhodophytina, two alternative positions placed just a single node away from each other.

Our results suggest that the secondary red endosymbiosis occurred either before the diversification of extant rhodophyte classes, or later, but from a so-far-unknown deep-branching lineage of red algae. This currently unknown lineage has either gone extinct (except for the complex plastids derived from it) or still exists, waiting to be discovered. Indeed, the discovery of a novel deep branching lineage in the phylogenetic vicinity of Rhodophyta would not be unprecedented (Gawryluk *et al*., 2019). Because smaller genetic divergence allows for better phylogenetic resolution, the discovery of a novel microbial lineage closely related to an organelle of interest may be crucial for resolving a disagreement between evolutionary scenarios, as demonstrated by the case of *Gloeomargaritales*, a recently described group of cyanobacteria (Couradeau *et al*., 2012). The close relationship of *Gloeomargaritales* to primary plastids has enabled resolving the controversy over whether these organelles derived from early- or late-branching cyanobacteria (Ponce-Toledo *et al*., 2017). Furthermore, if the sister position of cryptophyte NMs to Cyanidiophytina recovered in our analyses gets confirmed, it would support a freshwater and/or thermoacidophilic ancestry of CASH plastids, consistent with the notion that most of the early evolution of photosynthetic eukaryotes occurred in freshwater environments (Lewis 2017; Ponce-Toledo et al. 2017; Sánchez-Baracaldo et al. 2017).

## Acknowledgments

This work was funded by the European Research Council Advanced Grants Protist World (No. 322669, P.L.-G.) and Plast-Evol (No. 787904, D.M.) and the ERC Starting grant MacroEpik (No. 803151, L.E.). We thank the UNICELL single-cell genomics platform (deemteam.fr/en/unicell) for help in DNA preparation and Nanopore sequencing.

## Author contributions

LVFN, PLG, and DM designed and supervised the study. FvB and MC performed culturing, DNA extraction, and sequencing. LVFN, SAMG, and LE performed data collection and phylogenetic analyses. LVFN, SAMG, LE, PLG, and DM wrote and edited the manuscript. All authors read and approved of the final manuscript.

## Data Availability

All new sequences used in the phylogenomic analysis were deposited in NCBI GenBank under accession numbers PP113013–PP114032.

