## Supplementary figures and images for "Nucleomorph phylogenomics suggests a deep and ancient origin of cryptophyte plastids within Rhodophyta"

### Supplementary file S12

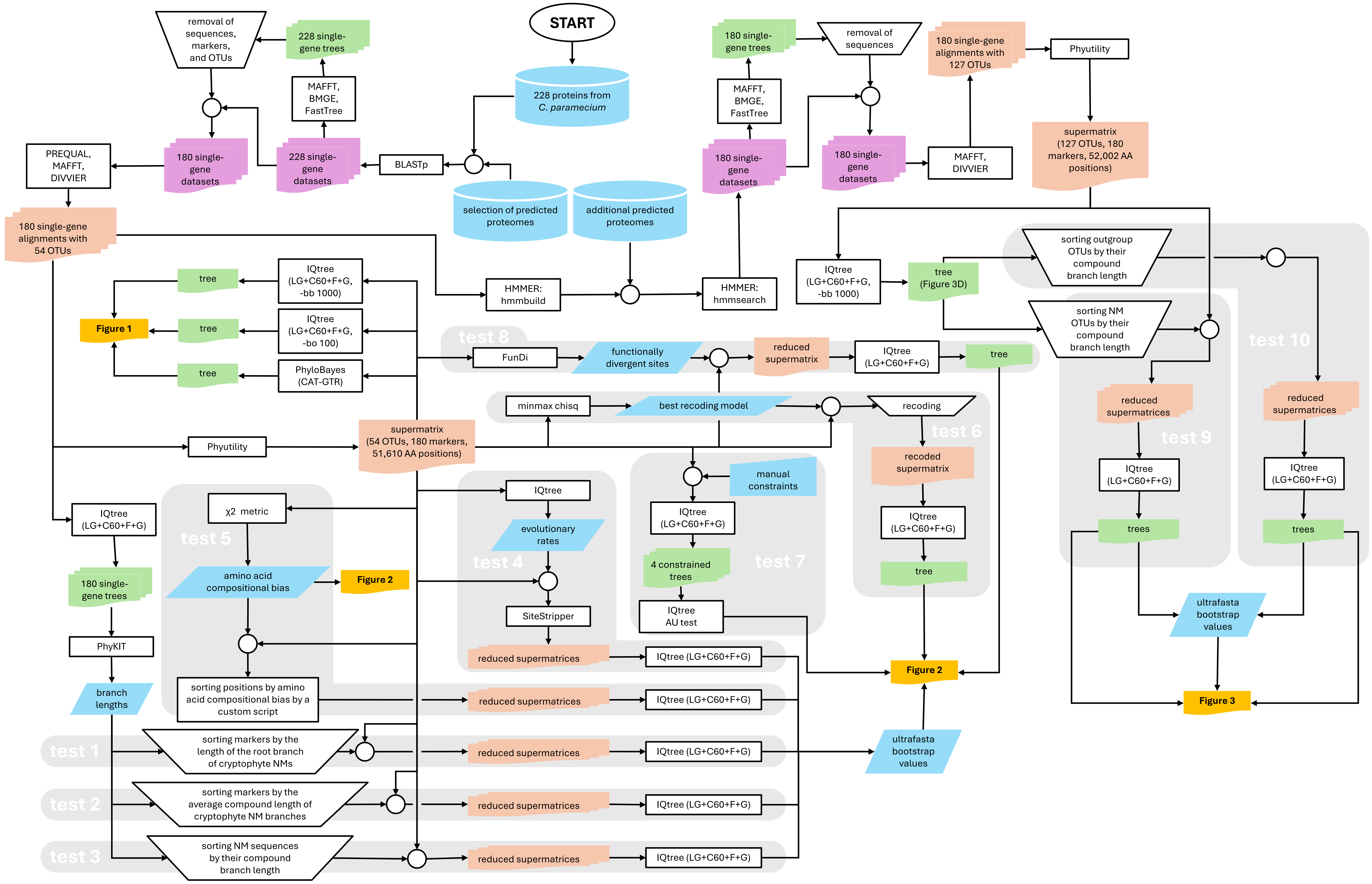
